# Machine-Learning prognostic models from the 2014-16 Ebola outbreak: data-harmonization challenges, validation strategies, and mHealth applications

**DOI:** 10.1101/294587

**Authors:** Andres Colubri, Mary-Anne Hartley, Mathew Siakor, Vanessa Wolfman, Tom Sesay, August Felix, Adam C. Levine, Pardis C. Sabeti

**Affiliations:** Harvard University, Department of Organismic and Evolutionary Biology, Cambridge, MA, USA; Broad Institute of MIT and Harvard, Cambridge, MA, USA; Howard Hughes Medical Institute, Chevy Chase, MD, USA; University of Lausanne, Faculty of Biology and Medicine, Switzerland; GOAL Global, Dublin, Ireland; International Medical Corps. Los Angeles, CA, USA; Sierra Leone Ministry of Health and Sanitation; Brown University, Warren Alpert School of Medicine. Providence, RI, USA; Harvard School of Public Health, Boston, MA, USA

**Keywords:** Ebola Virus Disease, Prognostic Models, Machine Learning, Data Visualization, Severity Score, mHealth, Supportive Care Guidelines, Clinical Intuition

## Abstract

**Background:** We created a family of prognostic models for Ebola virus disease from the largest dataset of EVD patients published to date. We incorporated these models into an app, “Ebola Care Guidelines”, that provides access to recommended, evidence-based supportive care guidelines and highlights the signs/symptoms with the largest contribution to prognosis.

**Methods:** We applied multivariate logistic regression on 470 patients admitted to five Ebola treatment units in Liberia and Sierra Leone during the 2014-16 outbreak. We validated the models with two independent datasets from Sierra Leone.

**Findings:** Viral load and age were the most important predictors of death. We generated a parsimonious model including viral load, age, body temperature, bleeding, jaundice, dyspnea, dysphagia, and referral time recorded at triage. We also constructed fallback models for when variables in the parsimonious model are unavailable. The performance of the parsimonious model approached the predictive power of observational wellness assessments by experienced health workers, with Area Under the Curve (AUC) ranging from 0.7 to 0.8 and overall accuracy of 64% to 74%.

**Interpretation:** Machine-learning models and mHealth tools have the potential for improving the standard of care in low-resource settings and emergency scenarios, but data incompleteness and lack of generalizable models are major obstacles. We showed how harmonization of multiple datasets yields prognostic models that can be validated across different cohorts. Similar performance between the parsimonious model and those incorporating expert wellness assessments suggests that clinically-guided machine learning approaches can recapitulate clinical expertise, and thus be useful when such expertise is unavailable. We also demonstrated with our guidelines app how integration of those models with mobile technologies enables deployable clinical management support tools that facilitate access to comprehensive bodies of medical knowledge.

**Funding:** Howard Hughes Medical Institute, US National Institutes of Health

## Research in context

### Evidence before this study

The recent Ebola virus disease (EVD) outbreaks have revealed the need for field-deployable tools that can be adapted to the heterogeneous spectrum of the disease across widely varying environments and resources. The 2015 report from the WHO’s Ebola Interim Assessment Panel addressed the shortcomings in the initial response to the 2014-2016 Ebola outbreak; it noted that “better information was needed to understand best practices in clinical management” and that “innovations in data collection should be introduced, including geospatial mapping, mHealth communications, and platforms for self-monitoring and reporting.” Even though there are detailed clinical guidelines available from WHO and other organizations, they are difficult to access by health workers in the field, since they are mostly available as documents in PDF format that are difficult to download and read on mobile phones. Machine learning prognostic models and mobile apps packaging these models and guidelines could be useful in making this knowledge more easily actionable, by quantifying the importance of each clinical feature in the prognosis of a particular patient, and thus help prioritizing the available interventions. We searched PubMed for studies using prognostic models and mHealth technology in the clinical management of EVD patients with the search terms “prognostic model”, “machine learning”, “mHealth”, “patient management” and “Ebola”. We only found one publication from van Griensven et al. in 2016 where the authors developed a prognostic model from a cohort of 85 patients, using electrolyte and metabolic abnormalities as predictors and testing it with a point-of-care device to stratify patients into risk groups.

### Added value of this study

The International Medical Corps (IMC) data used in this study includes 470 confirmed EVD cases from five different locations in Sierra Leone and Liberia. This is the largest and most diverse clinical EVD dataset available to date. This variety and sample size enabled us to construct a set of predictive models of moderate complexity using presentation information alone. The clinical and lab protocols were largely consistent across the five ETUs, making it possible to aggregate individuals into a single cohort after applying some data harmonization procedures. At the Sierra Leonean ETUs, the health workers also conducted observational wellness assessment of the patients, recorded at every daily round, and ranging from 0 (cured) to 5 (very sick patient). This data allowed us to create a set of prognostic models using different combinations of predictors, from a minimal model with only two predictors, age and viral load, to a parsimonious model incorporating several clinical features in addition to age and viral load. We were able to validate these models using two independent EVD patient cohorts from Sierra Leone, 106 treated at the Kenema Government Hospital and 158 treated in an emergency unit managed by GOAL Global. Our analysis showed that performance of the models was robust across the different datasets, and that the parsimonious model including the most detailed set of clinical and laboratory features had a performance that approximated the predictive power of the observational wellness assessments from health workers. We packaged all these models into a mobile app that offers evidence-based care guidelines compiled from WHO’s manuals for treatment and management of viral hemorrhagic fever patients. The models’ predictions are used by the app to highlight the interventions that are associated to the signs and symptoms with the largest contribution to patient death.

### Implications of all the available evidence

The construction of generalizable prognostic models requires datasets that are representative of entire the population under study. This is particularly challenging in low-resource settings or emergency situations, when data is often collected by different groups independently and without common definitions of clinical variables and protocols. In our analysis of the IMC dataset from Sierra Leone and Liberia, we were able to harmonize the records from 5 different sites thanks to the protocols put in place by IMC and shared across the sites. This enabled us to generate models that recapitulate prior findings regarding the clinical features at triage that are most predictive of death in EVD. Furthermore, the performance of models incorporating the observational wellness assessment is comparable or superior to the more detailed models including individual clinical features. This result suggests that machine learning approaches, when properly designed and implemented, and applied on rich-enough data, could approximate the clinical intuition or “gestalt” that physicians acquire through their experience in the field. Furthermore, the app is a first step in the development of a robust system that clinicians can use in the field. The prognosis predictions in the app are complemented with authoritative clinical care information provided by sources such as the WHO. In this way, the app could function both as a reference tool to improve training and adherence to protocol, as well as a support system that organizes clinical procedures more effectively around patient data.

## Introduction

The 2014-2016 outbreak of EVD caused a worldwide health crisis with more than 28,000 cases and 11,000 deaths, the vast majority of which occurred in the West African countries of Liberia, Sierra Leone, and Guinea. The recent outbreak in the Équateur Province of the Democratic Republic of the Congo (DRC) (1) and the subsequent, still ongoing, outbreak in the North Kivu Province (2) are evidence of the threat posed by EVD, even with the availability of experimental vaccines (3). Of particular concern, is the presence of outbreaks in regions with limited medical coverage such as the active conflict zone affected by the current outbreak.

Despite its notoriety as a deadly disease, the pathology of EVD includes a range of outcomes, spanning from asymptomatic infection to complex organ failure, with case fatality ratios (CFRs) of under 20% achievable in high-income countries where extensive resources can be applied on the few cases that were treated there. Clinical care of highly contagious diseases such as EVD in remote and low-resource settings is far more challenging, hindered by limited availability of trained personnel, restricted time that can be allocated to each patient due to difficult-to-wear personal protective equipment, and lack of supplies. Prioritizing time and material resources for high-risk patients is one approach to decrease overall mortality when subject to such constraints (4). A complementary approach is to use tools providing clinical instructions for management, training, and improved protocol adherence (5, 6).

We previously introduced the use of prognostic models that can be deployed on mobile applications (or apps for short) for the purpose of risk stratification in EVD (7). Prognostic models can enable the early identification and triage of high-risk patients, which could be useful in low-resource areas to better allocate supportive care. Health care workers could more frequently monitor those patients at increased risk and decide between standard and more aggressive therapy. Our original models were developed on the single publicly available dataset at that time by Schieffelin et al. (8), which includes 106 Ebola-positive patients at Kenema Government Hospital (KGH). These models outperformed simpler risk scores and allowed users to choose from various sets of predictors depending on the available clinical data. While this study showed the potential for such an approach, the models were limited by their geographical relevance (based on a single study site in one country, with a small patient cohort from one period during the outbreak). Furthermore, the prototype app in which the models were packaged was very simple, displaying only the severity score of the patient after the user entered the available clinical features and laboratory tests without further guidance.

We thus sought to create models with greatly expanded geographic relevance packaged in a new app that could provide risk-based guidance to health workers particularly in limited-resource settings. There are comprehensive materials available online, such as WHO and MSF’s clinical guidelines for viral hemorrhagic fevers, but these can be difficult to access by health workers in the field due to their book-like presentation, even if they are downloadable as digital files. This makes it hard finding relevant information quickly, tailoring it to the specific characteristics of the patients, or updating it as medical knowledge improves. Recently, a team of critical care and emergency medicine experts employed the Grading of Recommendations Assessment, Development, and Evaluation (GRADE) methodology to develop evidence-based guidelines for the delivery of supportive care to patients admitted to Ebola treatment units (9). These guidelines are particularly useful as a framework to organize the existing evidence, but still challenging to use for health workers in its current format as a research paper. Our goal is not only to make this information available through an app for health workers, but also tailor and organize care guidelines based on the severity score of the individual patients as predicted by validated prognostic models. In this way, the app could highlight recommendations and interventions that are most relevant given all the available information about the patient during triage.

Even though our models were derived from the largest and most diverse EVD patient cohort available to date (8, 10-14) -with 470 confirmed cases from Sierra Leone and Liberia-external validation across sites is critical to establish the geographic and demographic range to which the models may be generalized (15). Ideally, the models should be applied to data obtained independently from the cases originally used for model training, but even then, porting prognostic models from one center to another is challenging (16). To this end, we report two independent external validations on datasets collected at different health care centers with independent patient catchment areas on patients reporting at different time points of the epidemic. The first, includes the 106 Ebola-positive patients treated at KGH during the first months of the outbreak. The second, described by Hartley et al. (17), comprises 158 Ebola patients who were treated in an ETU run by GOAL global during the final months of the epidemic under conditions that should better represent future outbreak responses.

## Methods

### IMC Patient Cohort

The cohort used to develop the prognostic models in this study includes patient data collected at five ETUs operated by IMC in Liberia and Sierra Leone between September 15, 2014 and September 15, 2015. The ETUs were located at Lunsar (Port Loko District), Kambia (Kambia District), and Makeni (Bombali District) in Sierra Leone, and at Suakoko (Bong County) and Kakata (Margibi County) in Liberia. The majority of the patients did not come from holding units and presented directly to the IMC ETUs, with an overall Case Fatality Ratio (CFR) across the 5 ETUs of 58%. From the total 470 patients, 292 were treated at the three Sierra Leonean ETUs. Collection and archival protocols are detailed in Roshania et al (18). The Sierra Leone Ethics and Scientific Review Committee, the University of Liberia – Pacific Institute for Research & Evaluation Institutional Review Board, the Lifespan (Rhode Island Hospital) Institutional Review Board, and the Harvard Committee on the Use of Human Subjects provided ethical approval for this study and exemption from informed consent. A data sharing agreement was approved by IMC and the Broad Institute, following IMC’s Research Review Committee Guidelines (https://internationalmedicalcorps.org/document.doc?id=800).

### Data Collection

Trained nurses, physician assistants, physicians, and psychosocial support staff recorded patient demographic, clinical, and support data at least daily from admission to discharge on standardized paper forms – as part of routine clinical care and for epidemiologic purposes. The clinical and lab protocols were consistent across the five ETUs, making it possible to aggregate individuals into a single cohort, although some same variables were available in the Sierra Leonean and not in the Liberian ETUs, and vice versa. For example, only the three ETUs in Sierra Leone collected the wellness scale (WS) of the patients. WS is an observational assessment of patient wellness assigned by the physician or physician assistants, recorded at every daily round, and ranging from 0 (cured) to 5 (very sick patient), as described in Table 1. Local data officers entered this data into separate electronic databases at each ETU, which were combined together into a unified database. The RT-PCR data were obtained from four laboratories. The United States Naval Medical Research Center Mobile Laboratory in Bong County, Liberia, served the Bong and Margibi ETUs; the Public Health England (PHE) labs in Port Loko and Makeni in Sierra Leone processed samples from Lunsar and Makeni; the Nigerian Lab served the Kambia ETU. For the Liberian ETUs, RT-PCR was performed at the United States Naval Medical Research Center (NMRC) Mobile Laboratory in Bong County, while in Sierra Leone, the Public Health England (PHE) lab served the Lunsar and Makeni ETUs, and the Nigerian Lab served Kambia ETU. The lab assays and processing methodologies were similar across all these labs, with differences being the number of EBOV targets (NRMC used two targets, EBOV Zaire minor groove binding, PHE used only the EBOV Zaire target) and manual versus automated extraction. Further details on the lab methods are provided in the supplementary materials.

**Table 1.** Wellness scale. Interpretation of the 0-to-5 observational scale of patient wellness at the Sierra Leonean ETUs.

| *Wellness Scale* | *Interpretation* |
| --- | --- |
| 0 | Cured |
| 1 | Well: no symptoms: drinks and eats okay |
| 2 | Few symptoms: drinks and eats okay |
| 3 | Moderate symptoms: can walk, sit, and feed self |
| 4 | Sick: needs help to be fed, drink, and take medications |
| 5 | Very sick: needs IV fluids and medications, lots of help |

### Exploratory and Univariate Analysis

The primary variable of interest for patients admitted to the ETUs was final outcome (death or survival). The outcome of 5 patients was missing due to being transferred to another facility. The cycle threshold (CT) value is an inversely proportional proxy of viral load, with a cut-off of 40 cycles considered as negative. These values were calculated from PCRs performed on admission, or from the second PCR when the first was missing (performed no later than two days after admission and affecting 28 cases). Since non-survivor viremia is relatively stable (19), our estimation of risk for patients who ultimately died was probably minimally affected by this 48h delay. An overestimation of risk may be possible in survivors, as their viremia sharply decreases after the fifth day of illness, but this is situation is unlikely as most patients presented earlier (mean referral time was 4 days across all IMC sites). We carried out an initial univariate analysis of all factors against disposition, using the χ^2^ test with Yates correction for the binary variables, and the point biserial correlation test for numerical variables.

### Logistic Regression with Multiple Imputation

We constructed several logistic regression models to predict the binary outcome death/survival from the demographic, clinical, and laboratory data (viral load and malaria rapid test results) available at triage, using the R statistics software version 3.5.1. The main challenges in the modeling task were the high occurrence of missing values, and the lack of a pre-specified prognostic model. We handled missing data by initially discarding variables with more than 50% of missing values in either the Sierra Leonean or Liberian records. This criterion left some variables with up to 30% of missing values. Further removal of predictors, even those with high missingness, can increase bias in the regression coefficients (20), so we imputed the remaining missing values. Imputation methods require that the data is missing completely at random (MCAR), and we used the Little’s MCAR test, as implemented in the R package BaylorEdPsych (21). We applied multiple imputation with the aregImpute function from the R package Hmisc (22). This function generates a Bayesian predictive distribution from the known data, and outputs a number N of imputed datasets. Each missing value in the i^th^ imputation is predicted from an additive model fitted on a bootstrap sample with replacement from the original data. We set N=100, well above standard imputation guidelines (23). In order to construct the models, we first reviewed published research on the factors associated with death in EVD. We then complemented this prior knowledge with a variable selection procedure based on penalized logistic regression implemented with the R package Glmnet (24). We used an equal mixture of L1 and L2 penalties, also called Elastic Net regularization. The selection procedure consisted of fitting N times (one for each imputation) a fully saturated model including all variables, and then calculating the number of times the regression coefficient of each variable was greater than zero, thus indicating a positive association with death. We then applied the following inclusion criterion to define a non-redundant, parsimonious subset of predictors: we kept the variables that had a coefficient greater than zero in at least half of the penalized models. This criterion resulted in the inclusion of variables that were not as strongly correlated with outcome it the univariate analysis, such as bleeding and dyspnea, and elimination of variables that had a low P-value, like conjunctivitis. However, this would be anticipated as our variable selection procedure relied on the construction of multivariate models where marginal associations can become less significant when accounting for confounding dependencies. Once we identified a subset of variables in this manner, we constructed a family of non-penalized logistic regression models with the function lrm from the R package rms (25). This family includes a parsimonious model with all the variables obtained from the selection process, but also models that can be applied on smaller subsets of demographic information, clinical features and laboratory results, allowing us to use a less detailed model if not all variables are available at triage. This approach has been shown to outperform predictive value imputation, which consists of having only one full model and imputing missing values at prediction time using the data distribution from the training set (26). Each final model in the family was obtained by fitting N copies of the model on each imputed dataset, and then averaging those copies into a single model using the fit.mult.impute function in Hmisc. We conducted internal validation of the models using bootstrap resampling in order to obtain unbiased estimates of model performance-area under the curve (AUC), McFadden’s pseudo-R^2^, Brier score, accuracy, sensitivity and specificity-without decreasing sample size. A prediction was classified as death when the score from the model was over the 0.5 threshold. We calculated the confidence intervals (CI) of all the performance estimates using Fisher’s transformation (27), and converted logistic regression odds ratios (OR) to risk ratios (RRs) for the sign/symptoms present at triage using Zhang and Yu’s formula (28).

### External Validation

We did two external validations on independently collected datasets from Sierra Leone. The KGH dataset described by Schieffelin et al. (8) is the only such database to be made publicly available at the time of this study (https://dataverse.harvard.edu/dataverse/ebola). It includes 106 EVD-positive cases treated at KGH between May 25 and June 18, 2014. CFR among these patients was 73%. Sign and symptom data were obtained at time of presentation on 44 patients who were admitted and had a clinical chart. Viral load was determined in 58 cases. Both sign and symptom data and viral load were available for 32 cases. The GOAL dataset described by Hartley et al. (17, 29) includes 158 EVD-positive cases treated at the GOAL-Mathaska ETU in Port Loko between December 2014 and June 2015, where the CFR was 60%. Ebola-specific RT-PCR results and detailed sign and symptom data were available for all 158 patients. The Ebola-specific RT-PCRs recorded in the GOAL dataset were performed by the same PHE laboratory system as for the majority of the Sierra Leonean IMC data.

The KGH dataset includes RT-PCR data as viral load (VL) quantities expressed in copies/ml, but the corresponding CT values are no longer available. Since the IMC models use CT as a predictor, we transformed log(VL) to CT by solving for the standard qPCR curve transformation log(VL) = m×CT + c0, such that the minimum VL in the KGH dataset corresponds to the maximum CT in the IMC dataset, and vice versa. The assays used for diagnosing patients at KGH and IMC have very similar limits of detection (30–32), which justifies the methodology of our VL-to-CT transformation. We also note that a ≈10-fold increase in Ebola VL corresponds to a 3-point decrease in CT (33). Based on this relationship, −3/m in our formula should be close to 1, which is indeed the case (−3/m=0.976 using the m and c0 constants derived from the KGH and IMC data). Although different assays were used in the IMC ETUs, the log(VL) transformation remains unchanged if the curve is solved only with Liberian or Sierra Leonean CT data.

### Normalization of Cycle Threshold Values

We observed that CT values exhibits differences in their distributions between the Sierra Leonean and Liberian IMC ETUs, KGH, and GOAL ETU (Suppl. Figure S1). The mean and standard deviation of the CT values calculated over all cases are 21.82±5.16 (IMC ETUs in Sierra Leone), 27.67±5.45 (IMC ETUs in Liberia), KGH 26.05±6.00 (KGH), and 22.20±4.31 (GOAL ETU), with the differences being significant at P<0.0001. Essentially, the CT values form the IMC ETUs in Liberia and KGH are consistently higher in around 5 CT units for both fatal and surviving cases. Even though the RT-PCR assays and methods were comparable across sites, they were not identical (TaqMan in the Liberian ETUs, commercial Altona and in-house “Trombley” in the Sierra Leonean ETUs), therefore it is still possible they produced different CT values. Other possible reasons for the discrepancies in the CT distributions are unevenness in quality of care or in care-seeking behaviors, specially earlier during the outbreak. For example, it could be the case that patients in Liberia were less sick overall than patients in Sierra Leone, but received lower quality care, resulting fatalities at higher CT values than in Sierra Leone. This is possible, since most patients in Liberia were managed in the first couple months of IMC launching its response, before it was able to ramp up its human and material resources as well as perfect its treatment protocols. In order to account for those differences and to obtain a quantification of the viral load of the patients that is internally consistent for each site and comparable across sites, we normalized the CT values from each site/group of sites by subtracting the mean and dividing by the standard deviation. We used the normalized values to train and validate the models.

## Results

### Prognostic Potential and Prevalence of Signs and Symptoms Recorded at Triage

Triage symptoms reported by over 50% of fatal Ebola patients were anorexia/loss of appetite, fever, asthenia/weakness, musculoskeletal pain, headache and diarrhea (Table 2A). Few variables were significantly associated with patient outcome, suggesting that most clinical signs and symptoms have little predictive ability on their own, at least when considered at triage alone. Only CT, age (Table 2B), and jaundice (Table 2A) were associated with death at a level of P<0.05, while conjunctivitis, confusion, dyspnea, headache, and bleeding were weakly associated at P<0.15. However, statistical association of the variables when taken alone might be due to confounding effects in the data. Also, the prevalence of several triage symptoms was notably different between fatal and non-fatal outcomes, as can be seen by comparing their ranking (Suppl. Figure 2A) or their differential prevalence (Suppl. Figure 2B).

**Table 2.** Univariate analysis. Correlation between either binary (A) or continuous (B) clinical variables and the outcome of death. Marginal odds-ratios were obtained from the univariate logistic regression model for death using each variable alone as a predictor. For continuous variables, the Pearson’s R correlation coefficient is used and the odd-ratios correspond to inter-quartile range changes in the predictor. Notes: (*) variables not recorded at the Sierra Leonean ETUs, (**) variables not recorded at the Liberian ETUs.

| *Variable* | *Total (%)* | *Non-fatal (%)* | *Fatal (%)* | *Missing (%)* | *OR (95% CI)* | *P-value* |
| --- | --- | --- | --- | --- | --- | --- |
| *Jaundice* | 24/464 (5) | 4/197 (2) | 20/267 (7) | 1/470 (0) | 3.91 (1.31, 11.62) | 0.016 |
| *Conjunctivitis* | 128/464 (27) | 64/197 (32) | 64/267 (23) | 1/470 (0) | 0.66 (0.43, 0.99) | 0.054 |
| *Coma\** | 5/178 (2) | 0/83 (0) | 5/95 (5) | 292/470 (62) | NA | 0.096 |
| *Confusion\** | 16/178 (8) | 4/83 (4) | 12/95 (12) | 292/470 (62) | 2.86 (0.88, 9.23) | 0.120 |
| *Dyspnea* | 109/464 (23) | 39/197 (19) | 70/267 (26) | 1/470 (0) | 1.44 (0.92, 2.24) | 0.133 |
| *Headache* | 268/464 (57) | 122/197 (61) | 146/267 (54) | 1/470 (0) | 0.74 (0.51, 1.08) | 0.142 |
| *Bleeding* | 26/464 (5) | 7/197 (3) | 19/267 (7) | 1/470 (0) | 2.08 (0.86, 5.05) | 0.148 |
| *Asthenia/Weakness* | 334/464 (71) | 135/197 (68) | 199/267 (74) | 1/470 (0) | 1.34 (0.89, 2.02) | 0.187 |
| *Diarrhea* | 234/430 (54) | 96/187 (51) | 138/243 (56) | 35/470 (7) | 1.25 (0.85, 1.83) | 0.304 |
| *Malaria\*\** | 49/225 (21) | 17/94 (18) | 32/131 (24) | 241/470 (51) | 1.46 (0.76, 2.83) | 0.331 |
| *Dysphagia* | 112/464 (24) | 43/197 (21) | 69/267 (25) | 1/470 (0) | 1.25 (0.81, 1.93) | 0.374 |
| *Vomiting* | 197/464 (42) | 87/197 (44) | 110/267 (41) | 1/470 (0) | 0.89 (0.61, 1.29) | 0.587 |
| *Nausea\*\** | 94/286 (32) | 35/114 (30) | 59/172 (34) | 179/470 (38) | 1.18 (0.71, 1.96) | 0.613 |
| *Abdominal Pain* | 203/464 (43) | 89/197 (45) | 114/267 (42) | 1/470 (0) | 0.90 (0.62, 1.31) | 0.662 |
| *Bone/Muscle/Joint Pain* | 272/465 (58) | 118/197 (59) | 154/268 (57) | 0/470 (0) | 0.90 (0.62, 1.31) | 0.666 |
| *Throat Pain\** | 55/178 (30) | 24/83 (28) | 31/95 (32) | 292/470 (62) | 1.19 (0.63, 2.26) | 0.709 |
| *Cough\** | 61/178 (34) | 30/83 (36) | 31/95 (32) | 292/470 (62) | 0.86 (0.46, 1.59) | 0.738 |
| *Hiccups* | 55/464 (11) | 22/197 (11) | 33/267 (12) | 1/470 (0) | 1.12 (0.63, 1.99) | 0.805 |
| *Rash\** | 8/178 (4) | 3/83 (3) | 5/95 (5) | 292/470 (62) | 1.48 (0.34, 6.40) | 0.867 |
| *Chest Pain\** | 88/178 (49) | 41/83 (49) | 47/95 (49) | 292/470 (62) | 1.00 (0.56, 1.81) | 0.889 |
| *Photophobia\** | 24/178 (13) | 11/83 (13) | 13/95 (13) | 292/470 (62) | 1.04 (0.44, 2.46) | 0.892 |
| *Anorexia/*<br>*Loss of Appetite* | 316/464 (68) | 135/197 (68) | 181/267 (67) | 1/470 (0) | 0.97 (0.65, 1.44) | 0.946 |
| *Fever* | 349/464 (75) | 148/197 (75) | 201/267 (75) | 1/470 (0) | 1.01 (0.66, 1.54) | 0.944 |

**Table 2. Univariate analysis.** Correlation between either binary (A) or continuous (B) clinical variables and the outcome of death.
| *Variable* | *Mean non-fatal cases*<br>*(95% CI)* | *Mean fatal cases*<br>*(95% CI)* | *Missing*<br>*fraction (%)* | *Pearson's R* | *Odds-Ratio*<br>*(95% CI)* | *P-value* |
| --- | --- | --- | --- | --- | --- | --- |
| *Cycle Threshold* | 26.72 (15.92, 37.52) | 22.23 (11.18, 33.28) | 137/470 (29) | -0.37 | 0.33 (0.23, 0.47) | <0.0001 |
| *Wellness Scale\*\** | 2.49 (0.82, 4.17) | 3.20 (1.17, 5.22) | 247/470 (52) | 0.34 | 4.57 (2.44, 8.58) | <0.0001 |
| *Patient Age* | 28.49 (0.00, 58.72) | 32.03 (0.00, 72.10) | 4/470 (1) | 0.09 | 1.33 (1.01, 1.74) | 0.043 |
| *Body Temperature\*\** | 37.41 (35.50, 39.32) | 37.67 (35.34, 40.01) | 265/470 (56) | 0.12 | 1.39 (0.94, 2.06) | 0.099 |
| *Days since*<br>*Symptom Onset* | 4.28 (0.00, 12.03) | 4.20 (0.00, 9.90) | 108/470 (22) | -0.01 | 0.97 (0.76, 1.25) | 0.83 |

### Performance of Multivariate Logistic Regression Models

Our family of multivariate logistic regression models includes a parsimonious model with the most informative variables in the data. Several studies have previously identified single signs and symptoms statistically predictive for EVD mortality (17), such as high viral load (10, 11, 34, 35), hemorrhagic signs (10, 36, 37), confusion (11, 36, 38), extreme fatigue (38), asthenia (34, 38), and referral time (17). Taking this existing medical knowledge on EVD into consideration, and applying the variable selection procedure described in the methods, we obtained a set of variables including patient age as the only demographic parameter, the first available cycle threshold (CT) from PCR, and the clinical signs/symptoms bleeding, jaundice, dyspnea, dysphagia, and referral time (RT) recorded during triage. We also incorporated body temperature into the model, even though it was only recorded in the Sierra Leonean ETUs, because we considered that abnormally high and low temperatures could indicate either acute fever state or state of coma/shock. We used the categorical variable fever, available for all ETUs, as additional information to impute temperature for those subjects who were missing it. We applied restricted cubic splines to model the non-linear relationships between CFR and age (Suppl. Figure S2A) and CFR and body temperature (Suppl. Figure S2B). The actual values of age and temperature were the input of the spline fitting procedure, as implemented in the Hmisc package. The lack of a saddle in the CFR vs CT plot (Suppl. Figure S3) suggested that cubic splines were not required to model CT. Following Hartley’s (17) observation that viral load can act as a confounding factor for outcome and referral time, as very sick people with high viral load tend to present early and have higher mortality, we added a CT x RT interaction. Prior to applying multiple imputation, we found non-random patterns of missing values in the data. However, the Little’s MCAR test was satisfied at P=0.05 when considering the Sierra Leonean and Liberian records separately, meaning that the geographical origin was the main source of non-randomness. We still observed some weak non-random patterns in the Sierra Leonean records (P=0.03) for the RT variable, but they were caused by low values in CT (as seriously ill patients were less likely to report referral time). Inclusion of CT in the model controlled for this effect and ensured randomness within each subset of CT (high versus low).

In addition to the parsimonious model described above, we constructed an identical model but without body temperature as predictor, for comparison purposes and also to serve as a fallback when temperature is not recorded. We also constructed a clinical-only model that includes patient age, body temperature, and signs/symptoms at triage, which could be useful when PCR results are not available. We obtained the list of signs/symptoms in this model from application of the variable selection procedure after removing CT from the initial list of variables. This resulted in the inclusion of jaundice, bleeding, dyspnea, dysphagia, asthenia/weakness, and diarrhea. Finally, we constructed a minimal model only incorporating CT and age, which are the strongest predictors of outcome on their own, as observed in our data and reported by other researchers (14). Table 3 contains the validation indices and their 95% CIs of these models. Most indices are very similar between the parsimonious and minimal models, with largely overlapping CIs. The specificity and R^2^ are higher in the parsimonious models, while the sensitivity is slightly higher in the minimal model. The addition of temperature seems to consistently increase performance across all indices, but only by a minor amount. The clinical-only model shows consistently lower performance. Detailed description of all the models are provided in Suppl. Table S1.

**Table 3.** Validation indices for the prognostic models. These indices include UAC, McFadden’s pseudo R^2^ goodness-of-fit index, Brier score, overall accuracy, sensitivity, and specificity. The means and 95% confidence intervals were obtained with 200 iterations of bootstrap resampling.

|  | *Parsimonious*<br>*(95% CI)* | *Parsimonious*<br>*w/out temp.*<br>*(95% CI)* | *Clinical-only*<br>*(95% CI)* | *Minimal*<br>*(95% CI)* |
| --- | --- | --- | --- | --- |
| *AUC* | 0.75 (0.70, 0.79) | 0.74 (0.69, 0.78) | 0.64 (0.58, 0.69) | 0.76 (0.71, 0.80) |
| *R<sup>2</sup>* | 0.22 (0.17, 0.27) | 0.21 (0.16, 0.25) | 0.12 (0.09, 0.15) | 0.16 (0.13, 0.21) |
| *Brier* | 0.21 (0.16, 0.25) | 0.21 (0.16, 0.26) | 0.24 (0.19, 0.29) | 0.21 (0.16, 0.25) |
| *Accuracy* | 0.69 (0.64, 0.74) | 0.68 (0.63, 0.73) | 0.60 (0.54, 0.66) | 0.68 (0.63, 0.73) |
| *Sensitivity* | 0.80 (0.75, 0.84) | 0.79 (0.74, 0.83) | 0.70 (0.64, 0.75) | 0.81 (0.76, 0.85) |
| *Specificity* | 0.57 (0.51, 0.63) | 0.55 (0.49, 0.61) | 0.51 (0.44, 0.57) | 0.52 (0.46, 0.58) |

The calibration curves comparing the predicted and actual probabilities of death (Figure 1) suggest that the two parsimonious models represent the actual probabilities quite well, with some underestimation for low risk patients. The minimal and clinical-only and models exhibit a lower calibration, with the former underestimating the actual probabilities at both the low and high risk ends, and the later overestimating the actual probabilities for a wide range of risks above 60%.

**Figure 1.**
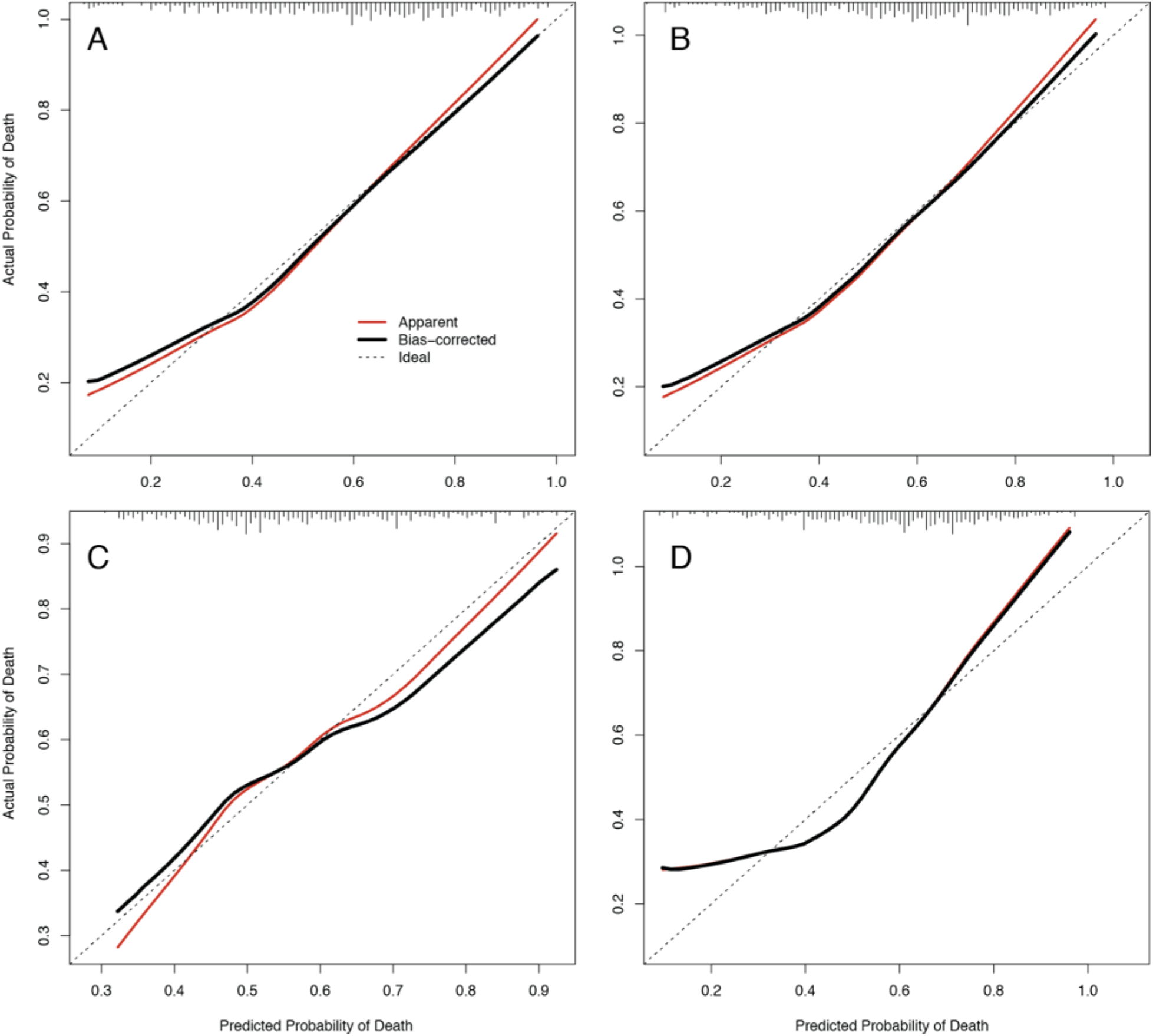
Bootstrap overfitting-corrected calibration curve. Estimated for the four prognostic models: parsimonious (A), parsimonious without body temperature (B), clinical-only (C), and minimal (D). Each plot contains the rug chart at the top showing the distribution of predicted risks. The dotted line represents the apparent calibration curve and the solid line shows the optimism-corrected calibration. A perfectly-calibrated model will fall along the diagonal. Generated with the calibrate function in the rms package.

The ranking of all of the variables by their importance in the parsimonious model, as measured by the Wald χ^2^ statistic, reveals that the most important variables are CT and patient age, with jaundice and bleeding coming in at a distant third and fourth place respectively, followed by body temperature and the CT-RT interaction (Figure 2A). The odds ratios (Figure 2B) indicate that presentation of either jaundice or bleeding are associated with more than a doubling of the odds of death, although their prevalence is low at 5% (Table 2). Transformation of the odds risks into risk ratios (Suppl. Table S1) yields a more reasonable estimate of 4% increase in risk associated to the occurrence of those symptoms at presentation. Finally, we compared the predictions from the parsimonious model when using the CT value from day 1 versus day 2, for those patients for whom both CT values where available. There were 9 such patients (5 fatal cases), and the results were largely consistent (Supp. Table S2) with the predictions using CT from day 1 misclassifying only one case as fatal, and those using CT from day 2 also misclassifying second case as surviving. The predicted scores as very similar between the two sets of predictions, with only the second case showing a markedly different score (0.87 vs 0.24) due to the fact that the CT of this subject increased dramatically from 17 to 33 in the second day at the ETU.

**Figure 2.**
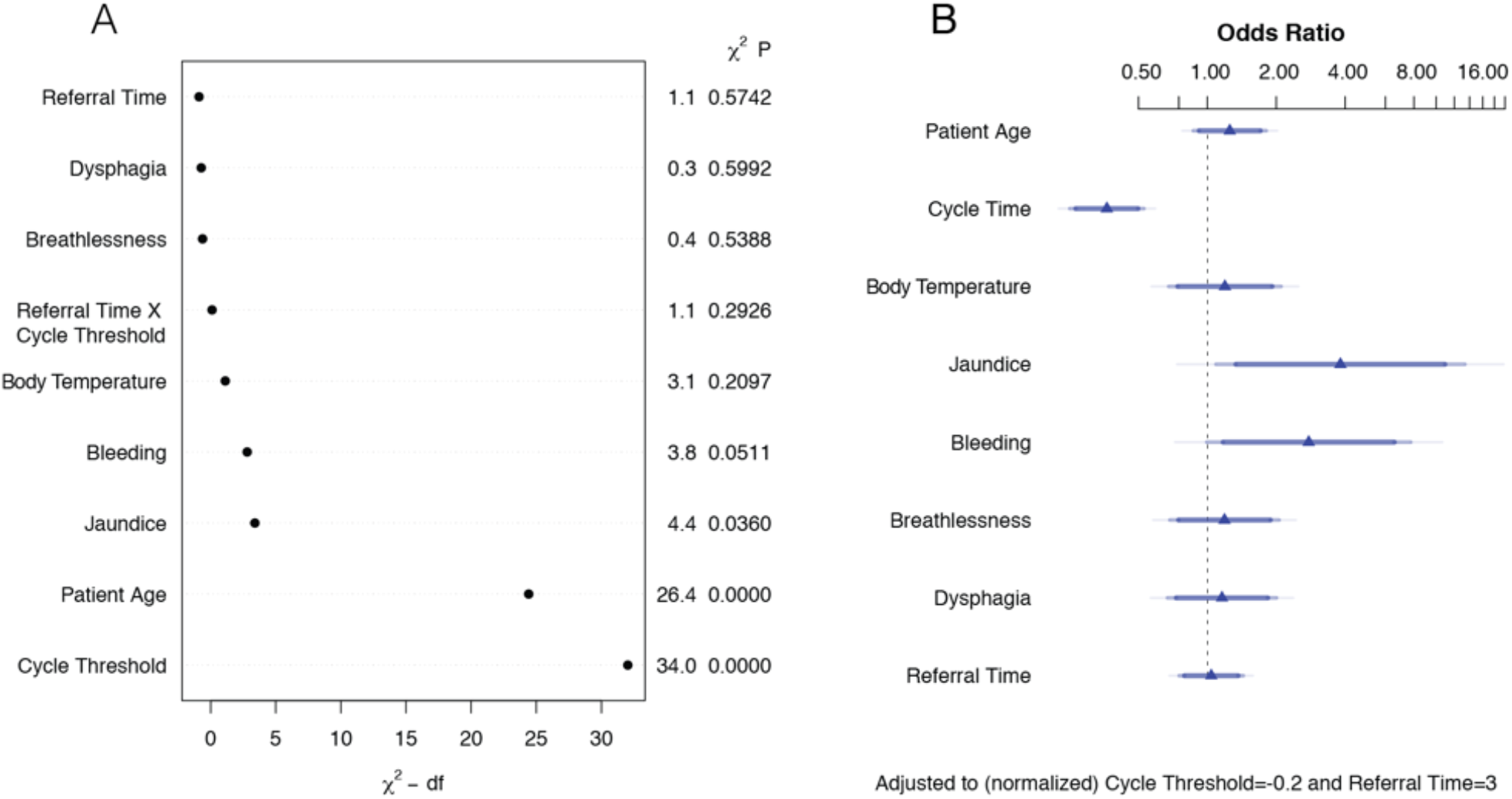
Evaluation of predictor variables in the parsimonious model. Ranking of the variables according to their predictive importance in the model, as measured by the χ^2^-d.f. (degrees of freedom) statistic (A). Odds ratios for all the variables, using interquartile-range odds ratios for continuous features, and simple odds ratios for categorical features (B). Generated with the anova.rms and summary functions in the rms and base packages in R.

### External validation

External validation on the 158 EVD-positive patients in the GOAL dataset shows that the four models described earlier improve their performance with respect to the internal validation. CT was missing in 37 patients, and both CT and RT were missing in 53, so the parsimonious models were validated on 105 patients and the minimal model on 121. The clinical-only model was validated on all the 158 patients. We obtained AUCs of 0.82, 0.84, 0.73, and 0.82 for the parsimonious, parsimonious without temperature, clinical-only, and minimal (Table 4). In terms of the accuracy, sensitivity, and specificity, all the models, with the exception of the clinical-only, performed similarly well with accuracies around 74%. Excluding again the clinical-only model, sensitivity ranged between 78% and 82%, while specificity was lowest for the minimal model at 61% and highest for the parsimonious model at 68%. Therefore, it is not possible to pick an overall best model between the three including CT, but depending on what is the priority in the predictive task, higher sensitivity or specificity, one could favor either the parsimonious or the minimal.

**Table 4.**
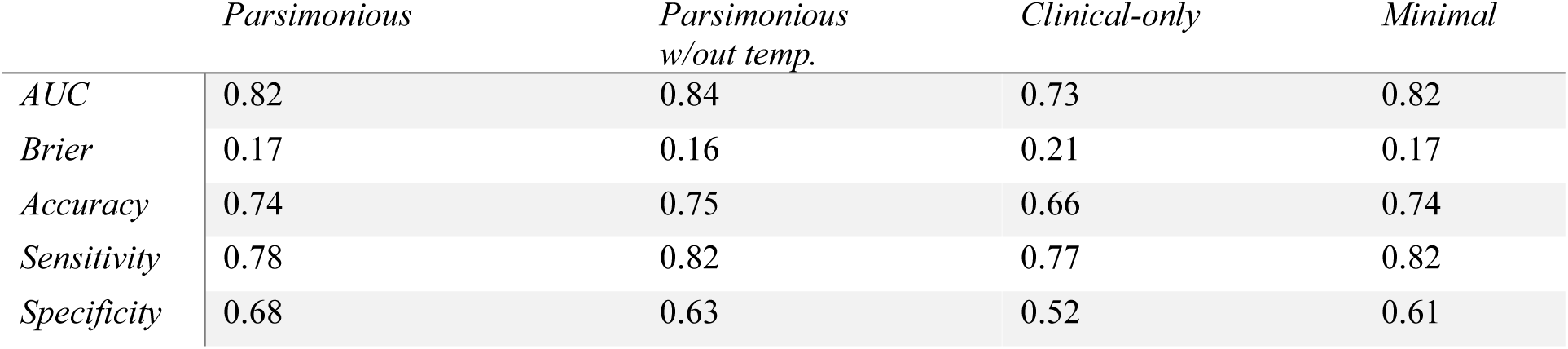
External validation on the GOAL dataset. The AUC, Brier, accuracy, sensitivity, and specificity indices were calculated on all the records from the GOAL dataset with complete data.

|  | *Parsimonious* | *Parsimonious*<br>*w/out temp.* | *Clinical-only* | *Minimal* |
| --- | --- | --- | --- | --- |
| *AUC* | 0.82 | 0.84 | 0.73 | 0.82 |
| *Brier* | 0.17 | 0.16 | 0.21 | 0.17 |
| *Accuracy* | 0.74 | 0.75 | 0.66 | 0.74 |
| *Sensitivity* | 0.78 | 0.82 | 0.77 | 0.82 |
| *Specificity* | 0.68 | 0.63 | 0.52 | 0.61 |

External validation on the KGH dataset is shown in Table 5. Only 11 patients had enough complete records to be used for validation of the parsimonious models, so the missing values were imputed using the same multiple imputation approach described in the methods. In this way, the two parsimonious models could be applied on all the 106 KGH cases. As it was the case with the GOAL validation, all models with the exception of the clinical-only exhibit consistent performance in terms of AUC and overall accuracy, with analogous differences in terms of specificity and sensitivity. The parsimonious models have higher specificity, while the minimal is more sensitive. The variation due to the imputation is minor and does not changes these results.

**Table 5.** External validation on the KGH dataset. The AUC, Brier, accuracy, sensitivity, and specificity indices were calculated on all the records from the KGH dataset that had enough data to evaluate the clinical-only and minimal models. The results for the parsimonious models were obtained after imputing missing values in the KGH patients, and the performance indices include the mean and standard deviation over 100 multiple imputations.

|  | *Parsimonious*<br>*(SD)* | *Parsimonious*<br>*w/out temp. (SD)* | *Clinical-only* | *Minimal* |
| --- | --- | --- | --- | --- |
| *AUC* | 0.78 (0.04) | 0.78 (0.04) | 0.72 | 0.82 |
| *Brier* | 0.19 (0.01) | 0.18 (0.01) | 0.22 | 0.17 |
| *Accuracy* | 0.73 (0.03) | 0.72 (0.03) | 0.59 | 0.74 |
| *Sensitivity* | 0.73 (0.03) | 0.73 (0.03) | 0.56 | 0.82 |
| *Specificity* | 0.88 (0.03) | 0.86 (0.02) | 0.75 | 0.61 |

### Wellness Scale Models

We constructed four additional models incorporating the wellness scale (WS) variable in place of the detailed clinical signs and symptoms used in the previous models: wellness parsimonious (including CT, patient age, body temperature, WS, RT and RTxCT), wellness parsimonious without temperature, wellness clinical-only (including patient age, body temperature, and WS), and wellness minimal (including CT, patient age, and WS). The performance indices of these models, obtained from internal validation on the Sierra Leonean patients from the IMC dataset, are shown in Table 6.

**Table 6.** Validation of the wellness scale models. These models were evaluated using the same indices of performance as the previous models: AUC, R^2^, Brier, accuracy, sensitivity, and specificity. The means and 95% confidence intervals were obtained with 200 iterations of bootstrap resampling.

|  | *Wellness parsimonious*<br>*(95% CI)* | *Wellness parsimonious*<br>*w/out temp. (95% CI)* | *Wellness clinical-only*<br>*(95% CI)* | *Wellness minimal*<br>*(95% CI)* |
| --- | --- | --- | --- | --- |
| *AUC* | 0.80 (0.74, 0.84) | 0.81 (0.76, 0.85) | 0.74 (0.68, 0.80) | 0.80 (0.74, 0.84) |
| *R<sup>2</sup>* | 0.31 (0.24, 0.38) | 0.30 (0.23, 0.37) | 0.19 (0.14, 0.25) | 0.25 (0.19, 0.32) |
| *Brier* | 0.19 (0.13, 0.24) | 0.18 (0.13, 0.23) | 0.20 (0.15, 0.26) | 0.19 (0.13, 0.24) |
| *Accuracy* | 0.73 (0.66, 0.79) | 0.73 (0.67, 0.79) | 0.69 (0.62, 0.75) | 0.72 (0.65, 0.77) |
| *Sensitivity* | 0.82 (0.76, 0.86) | 0.82 (0.77, 0.86) | 0.79 (0.72, 0.83) | 0.81 (0.75, 0.85) |
| *Specificity* | 0.63 (0.55, 0.70) | 0.63 (0.55, 0.70) | 0.57 (0.48, 0.64) | 0.61 (0.53, 0.68) |

All the performance indices of these new models are higher than the corresponding original models. In fact, the performance of the worst performing model in this set, the wellness clinical-only, is comparable to that of the parsimonious model including the detailed signs and symptoms. The calibration curves for the parsimonious models with and without wellness scale are almost indistinguishable, but it is possible to note an improvement for the clinical-only and minimal models with the addition of the wellness scale (Figure 3).

**Figure 3.**
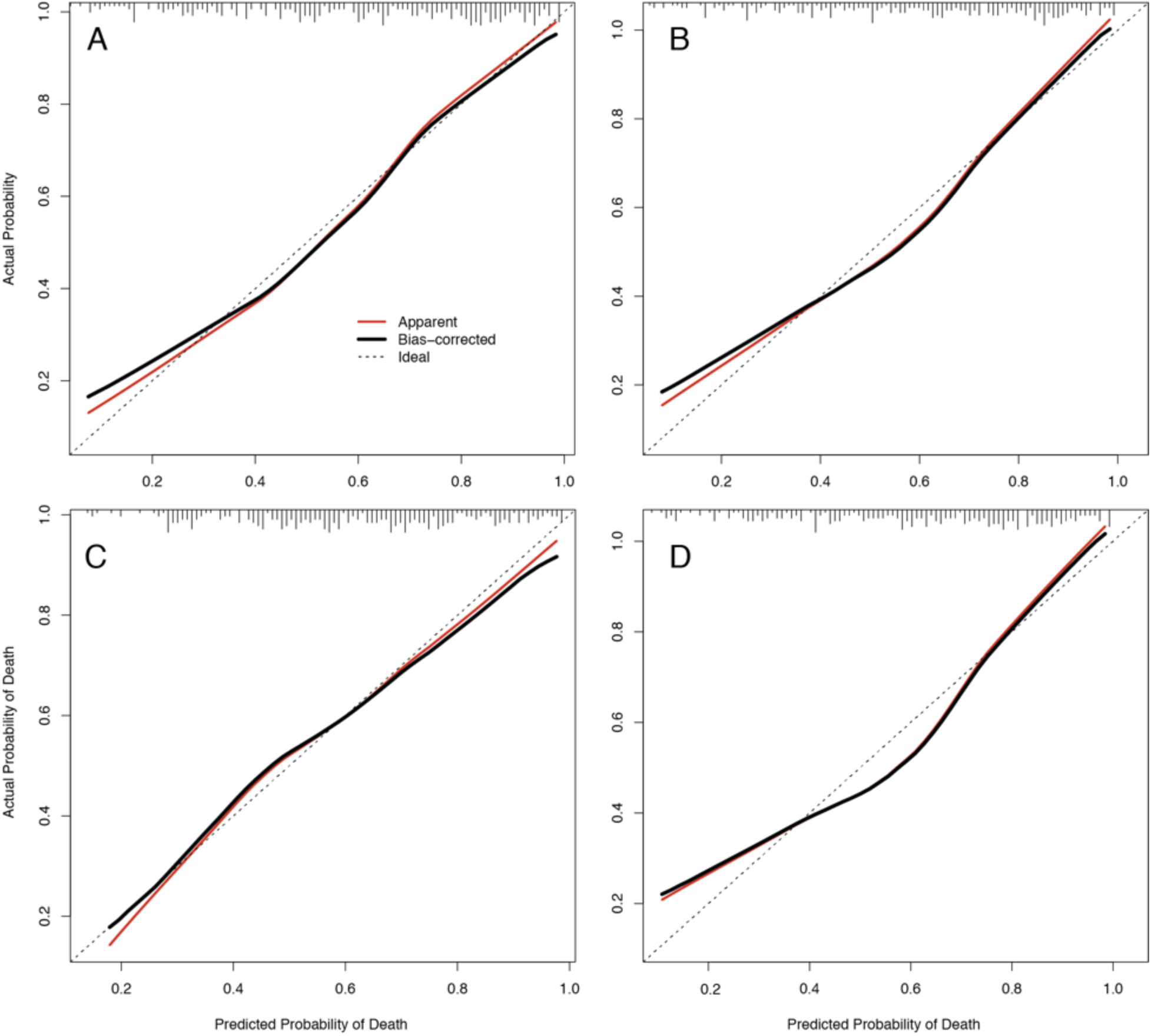
Bootstrap overfitting-corrected calibration curves for the wellness models. Estimated for the four prognostic models using the wellness scale as predictor: wellness parsimonious (A), wellness parsimonious without body temperature (B), wellness clinical-only (C), and wellness minimal (D). Each plot contains the rug chart at the top showing the distribution of predicted risks. The dotted line represents the apparent calibration curve and the solid line shows the optimism-corrected calibration. A perfectly-calibrated model will fall along the diagonal. Generated with the calibrate function in the rms package.

The wellness assessments were available for 223 of the 292 patients treated at the three ETUs in Sierra Leone, and so the WS value was imputed for the 89 Sierra Leonean patients without it. To evaluate the effect of imputation in the models, we fitted the four wellness models only on those patients with known WS. The performance of the refitted models (Suppl. Table S3) is very close to that of the models derived from imputed data, which indicates that variance inflation caused by the imputation is minimal.

### Ebola Care Guidelines App

We created a multilingual Android app for health workers, named Ebola Care Guidelines, that integrates available patient care and management guidelines for EVD patients with the prognostic models described earlier. The app only requires internet connectivity to be installed the first time, and it can be used even when the device is offline afterwards. This is an important feature as emergency health workers are often deployed in rural or remote locations with limited internet access. The home screen of the app shows a list of supportive care recommendations for Ebola fever patients, compiled from Lamontagne et al. (9), and WHO and MSF’s care and management guidelines for hemorrhagic fevers (Figure 4A). The list is categorized by intervention type (such as oral rehydration, parenteral administration of fluids, monitoring of viral signs and volume status, etc.). Selecting a guideline from this list provides a summary description and specific interventions related to that guideline (Figure 4B), obtained from WHO’s manual for the care and management of patients in Ebola Care Units/Community Care Centers (39) and MSF’s clinical management of patients with viral hemorrhagic fever pocket guide for front-line health workers (40). When users select a specific intervention, the app redirects to the corresponding page in the document (Figure 4C). Accessing this information does not require entering any patient data; in this way the app can be useful simply as a targeted entry point to these detailed guidelines. Users can also input patient information (age, sex, pregnancy status, weight, height), clinical signs and symptoms recorded at triage, laboratory results, and the wellness scale (WS) from the first clinical rounds after admission (Figure 4D), through a built-in form, or via a separate CommCare (https://www.commcarehq.org) app. After all, or some, of this data is recorded, the app computes the severity score of the patient by selecting the appropriate model for the available data. The app offers a visualization of the score and the magnitude of patient-specific contributions for each feature included in their score, where the score value is shown at the top in a color-graded scale and patient-specific feature contributions to that score are depicted in a bar chart summary page (Figure 4E). Each clinical feature is linked to one or more care guidelines, so that when that feature is present in the data, the corresponding guidelines are highlighted (Figure 4F). The total severity score can also be linked to specific guidelines when it is over a threshold defined in the app’s settings. Those guidelines will also be highlighted when the score is higher than a user-defined threshold. This feature is designed to bring the user’s attention to the recommendations that could be most relevant given the clinical manifestation of the patient. It is important to note that the app does not make any choices for the user that deviates from the guidelines, it simply directs the user to the section in the guidelines that advises them in that decision. The trigger signs/symptoms linking to the guidelines are straightforward, since are precisely those listed in the guidelines themselves (e.g.: vomiting and diarrhea for the guidelines on ORS/IV fluids). The app’s source code is available on GitHub under the MIT license (https://github.com/broadinstitute/ebola-care-guidelines), where instructions on how to update the models and the guidelines included in the app are provided as well.

**Figure 4.**
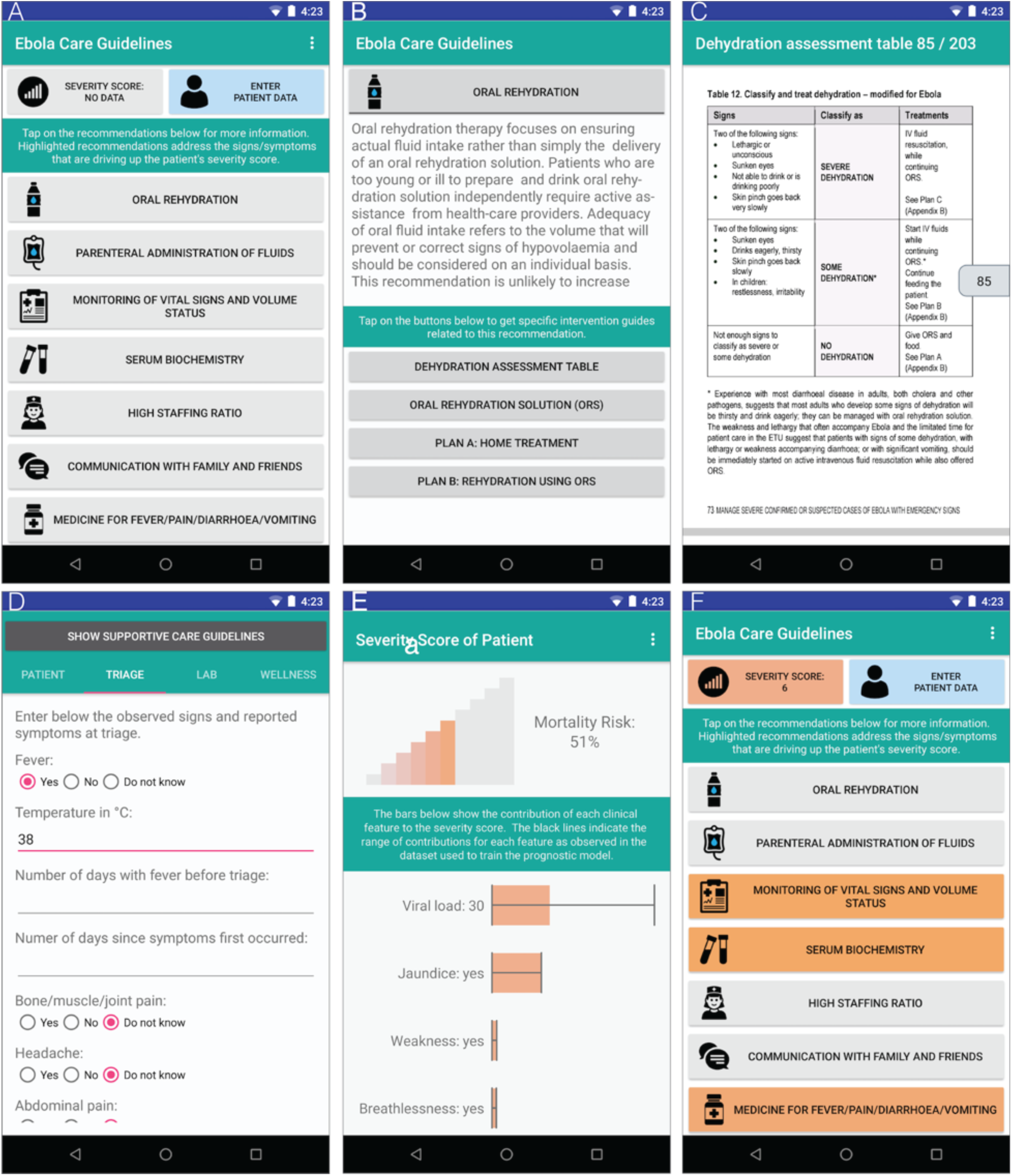
Ebola Care Guidelines app. The home screen presents the list of recommendations (A), which can be selected to access specific interventions associated to each recommendation (B). Selecting a specific intervention or guideline redirects the user to the corresponding section in the WHO’s manuals for care and management of hemorrhagic fever patients (C). The app allows the users to enter basic demographic information (age, weight), vitals, signs & symptoms at presentation, lab data (CT value from first RT-PCR and malaria test result), and wellness scale (D). Based on the available data, the app calculates the severity score of the patient using the suitable prognostic model and presents a customized risk visualization (E). The recommendations that are associated with the presentation signs and symptoms are highlighted in the home screen (E).

## Discussion

### IMC Models Recapitulate Prior Findings and Highlight the Importance of Clinical Intuition

The purpose of this study was two-fold: first, to present generalizable machine learning prognostic models for EVD derived from the largest multi-center clinical dataset available to date, externally validated across diverse sites representing various periods of the largest Ebola epidemic on record; and second, to show how these models could guide clinical decisions by organizing existing knowledge of patient care and management more efficiently and making it easily available as a mobile app. The discriminative capacity of the IMC models is robust across the training set and two independent validation sets, which were obtained at distinct times during the epidemic, with AUCs ranging from 0.75 up to 0.8. Furthermore, these models recapitulate several findings reported earlier in the literature and also reveal further associations between mortality and clinical signs/symptoms. The most informative descriptors for predicting EVD outcome are patient age and viral load, and models incorporating just these two variables exhibit good performance. Occurrence of jaundice or bleeding at initial presentation are important predictors of death, but both have low incidence at triage among the patients in the IMC cohort of only 5%. In contrast, more widespread EVD manifestations such as dyspnea, dysphagia, and weakness have a much weaker correlation with mortality, at least based on their presence at triage, which seems to suggest that presentation of these clinical features says little about the clinical evolution of the patient. Even though fever is a non-specific symptom with no predictive power, body temperature at triage is informative by increasing specificity in the predictions.

The performance of models incorporating the clinical wellness score is at least comparable or superior to the more detailed models including individual clinical features deemed as the most predictive in our variable selection process. This is in fact a very important result in our study, since it suggests that machine learning approaches, when properly designed and implemented, and applied on rich-enough data, could approximate the clinical intuition that physicians acquire through their experience in the field. Conversely, this also provides evidence that good bedside intuition can very effectively integrate the individual indicators of the patient’s clinical status into an overall health assessment of great predictive power. These models may be useful in emergency situations when the appropriate experience is unavailable or under-developed.

Our prognostic models and guidelines app can be easily updated with new data. With increasing knowledge on the management of Ebola, we expect the mortality rate to decrease over time or to be more reliably estimated by earlier symptoms. New information from the ongoing outbreak in DRC or future epidemics would enable us to train and update the models to more appropriate calculations that are adapted to improving standards of care, and geographically divergent variables. For example: health-care seeking behavior in the conflict-afflicted DRC would be undoubtedly more limited than that in West Africa in 2014-16, and perhaps other factors will be more predictive of death. So not only are the models adapted to the newly available information but the new models automatically link up to the easily-updatable guidelines as they become available.

### Data-harmonization Challenges

In order to account for different levels of clinical detail collected at the ETUs, we constructed a family of prognostic models that range from models requiring only clinical signs/symptoms or age and viral load, to more complex models incorporating a mixture of laboratory data, signs/symptoms, and even observational assessments from experienced health providers. We harmonized the available data to extract the most value from the IMC cohort. For example, in order to aggregate CT values from different PCR labs, we applied an intra-site normalization to ensure that values were comparable across sites even if different assays produced somewhat different CT values. We had to discard several potentially informative clinical variables such as confusion and coma since they were not available in all the IMC ETUs. We were able to incorporate body temperature into the models, even though it was not recorded in the Liberian ETUs, by using multiple imputation with fever as an auxiliary variable. Increased specificity in the validation sets of the models incorporating temperature justified this modeling decision.

### Validation Strategies

Our prognostic models, particularly those incorporating the parsimonious sets of predictors, perform well on two independent datasets used for external validation. These datasets have a wide temporal, geographic and clinical scope. A major difference between these datasets was the time during which they were collected, with the KGH data representing an earlier time point, with less refined treatment protocols, increased patient volume and admission intensity with a larger number of patients delayed during transfers from holding centers. On the other hand, the GOAL dataset includes patients from the final months of the epidemic with a 13% lower CFR. Thus, as may be expected, the models underestimated the observed risk for patients of the KGH cohort, while observed risk was slightly overestimated in the GOAL cohort. The IMC training dataset covers a much broader temporal window of the epidemic as well as a wider catchment area, spanning several districts across two countries, which may explain its consistent performance in these disparate populations.

### mHealth Applications

Finally, with the Ebola Care Guidelines app we aimed at developing a robust and flexible mHealth system that clinicians can trust in the field and in emergency situations. An initial step in that direction is to complement the prognosis predictions with authoritative clinical care information provided by sources such as the WHO. In this way, we envision the app both as a reference tool to improve training and adherence to protocol, as well as a support system that organizes clinical procedures more effectively around the patient’s data. The integration of mHealth platforms with rapid point of care diagnostic kits (41, 42) has the potential to realize the concept of a “pocket lab” (43), which could be used outside laboratory settings and during health emergencies. The ultimate goal of these platforms is to aid clinical management decisions on the ground by enabling the design of clinical support systems for front-line workers that better organize and provide access to the existing medical knowledge on viral hemorrhagic fevers, tailored to reflect the patient’s clinical signs, symptoms and laboratory results. Our approach is generalizable in the sense that it can be applied to create new mobile apps for other tropical diseases affecting rural and low-resource areas, and also provides a mechanism to keep the clinical guideline apps updated as the medical knowledge is refined, and more accurate prognostic models are developed in the light of new and better data. The current update process simply involves bundling the specifications of the new models (predictor variables and coefficients), rebuilding the app, and releasing it through Android’s app store. Future versions of the app would not even require to be updated, as they could retrieve new models directly from a remote server. We believe that if clinical staff can obtain actionable information from these data-derived tools, then they may be incentivized to generate more and higher-quality data, which could then be incorporated back into the models, creating a positive feedback loop which drives increased precision. Further, the visualization of the clinical make up of prediction models (such the charts provided in our app) provides a learning platform that builds informed clinical experience rather than simply replacing it.

### Limitations

Despite being the largest EVD prognosis modeling study to date, the amount and quality of available clinical data is still limited. We accounted for these limitations by harmonizing the data from different ETUs and applying various statistical techniques recommended for prognosis modeling (multiple imputation, bootstrap sampling, external validation). Even with the help of these approaches, data might be affected by variations in clinical assessments from clinicians with varying levels of experience, errors in data collection (including patient symptom recall or history taking skills), and differences in lab protocols. Ultimately, future predictive models will require larger and better datasets, and consensus mechanisms to ensure that data is consistent across sites.

### Conclusions

It is possible to generate generalizable machine learning prognostic models if data from representative cohorts can be harmonized and validated properly. The use of low-cost mHealth tools on the ground incorporating the insights gained from such models, in combination with effective data collection and sharing among all stakeholders, will be key elements in the early detection and containment of future outbreaks of Ebola and other emerging infectious diseases.

## Availability of source code, data, and app

The source code of all the modeling steps, from parameter fitting to internal and external validation, is available as a fully documented Jupyter notebook, deposited online at https://github.com/broadinstitute/ebola-imc-code. The source code of the Ebola Clinical Guidelines app is available at https://github.com/broadinstitute/ebola-care-guidelines. Refer to IMC’s Ebola Response page (https://internationalmedicalcorps.org/ebola-response), for instructions on how external researchers can access the data. The app is freely available on Google Play: https://play.google.com/store/apps/details?id=org.broadinstitute.ebola_care_guidelines

## Acknowledgments

We would like to thank the governments of Liberia, Sierra Leone, and Guinea for contributing to International Medical Corps’ humanitarian response. We would also like to thank all of our generous institutional, corporate, foundation, and individual donors who placed their confidence and trust in International Medical Corps and made our work during the Ebola epidemic possible. We would also like to thank the United States Naval Medical Research Center, Public Health England, the European Union Mobile Laboratory, and the Nigerian Laboratory for providing laboratory data to our Ebola Treatment Units. We would like to further acknowledge all members of our Research Review Committee and other technical teams that contributed to this research. Finally, we would also like to thank our clinical, WASH, and psychosocial teams as well as all of our monitoring and evaluation staff, including the data collection officers at each of our ETUs. AC would like to thank Mary Lynn Baniecki and Christian Matranga for insightful discussions on the EBOV qPCR assays, and Christopher Moxon for critical feedback on the manuscript. Finally, we thank the patients included in this study whose data has and will continue to make invaluable contribution to improving future Ebola care.

