## Supplementary figures and tables for "Machine-Learning prognostic models from the 2014-16 Ebola outbreak: data-harmonization challenges, validation strategies, and mHealth applications"

### Supplementary materials

#### Figure S1: Distribution of RT-PCR Cycle Threshold (CT) across all sites

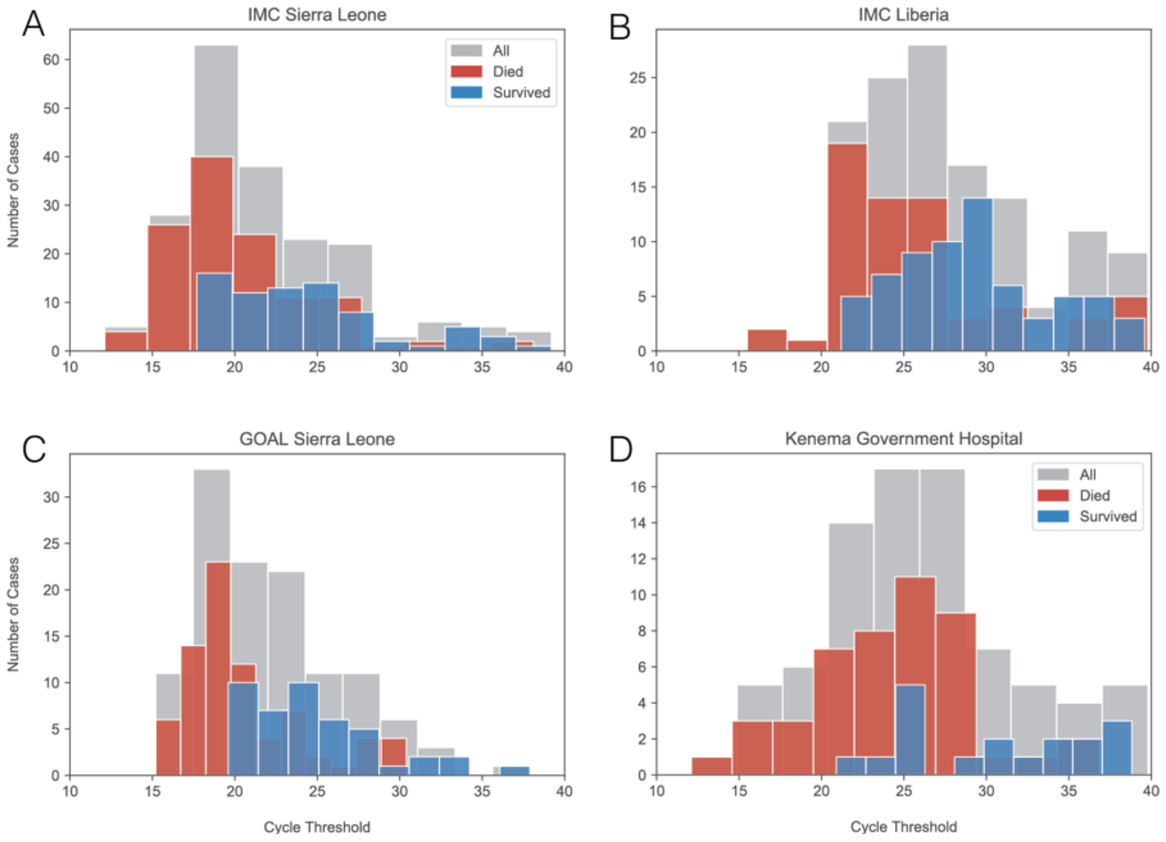


*The histograms represent the distributions of the CT values from all the sites considered for model training and validation:(A) ETUs operated by IMC in Sierra Leona, located at Lunsar (Port Loko District), Kambia (Kambia District), and Makeni (Bombali District). (B) ETUs operated by IMC Liberia, located at Suakoko (Bong County) and Kakata (Margibi County). (C) GOAL-Mathaska ETU in Port Loko, Sierra Leone. (D) Kenema Government Hospital in Sierra Leone. In each plot, the grey histogram includes all cases, the red histogram only the fatal cases, and the blue histogram only the survivors.*

**Figure S2: Prevalence of clinical signs and symptoms recorded at triage**

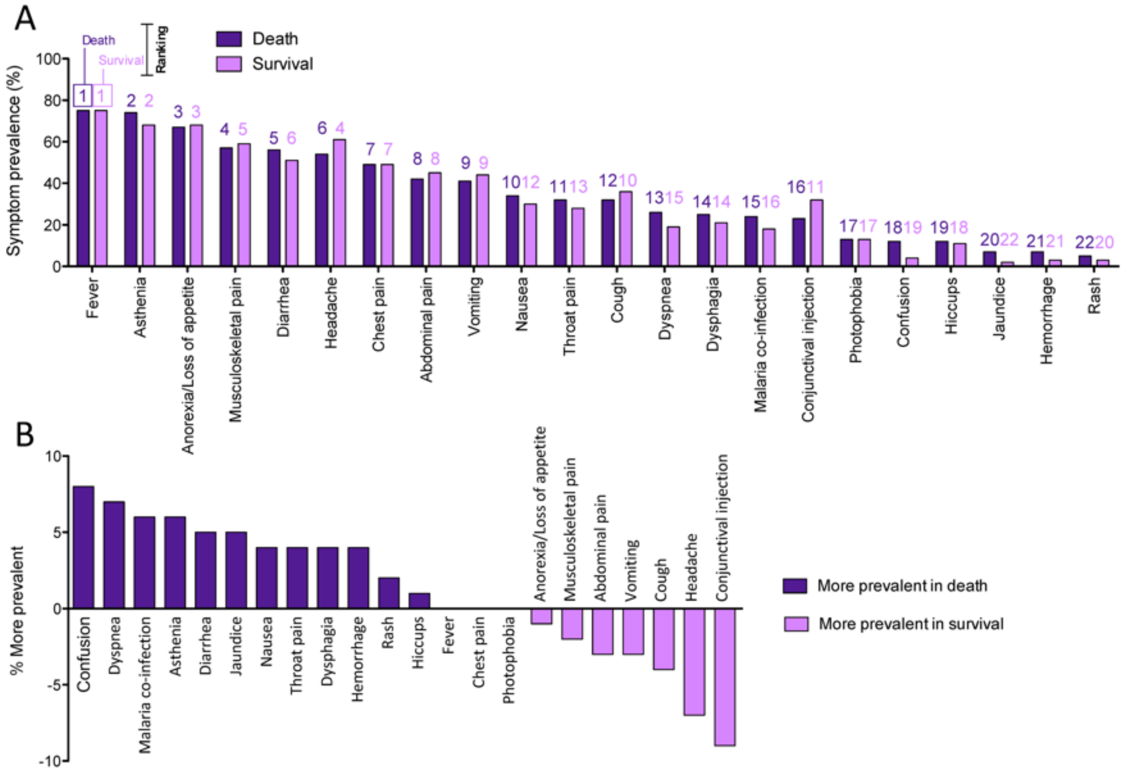


*(A) Prevalence of clinical characteristics at triage amongst Ebola patients who either survived or died, ranked according to the prevalence in fatal outcomes. Rankings from 1–22 are listed above each bar: purple for the outcome of death and pink for survival. (B) Differences in symptom prevalence between EVD survivors and those who died. Positive values are more prevalent in fatal outcomes. Negative values are more prevalent in survivors.*

**Figure S3: Case fatality rate as a function of patient age and body temperature**

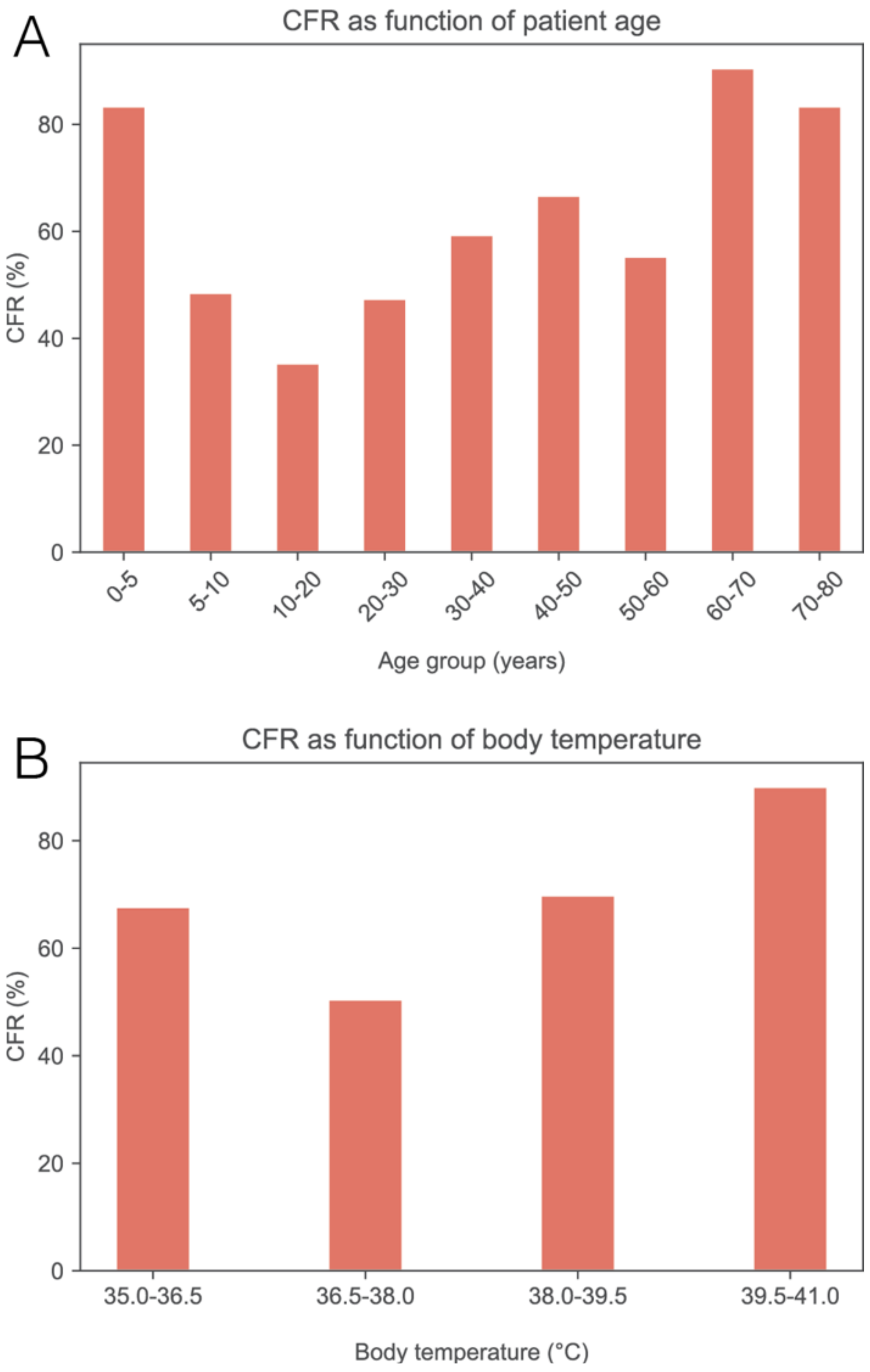


*Case Fatality Rate (CFR) as a function of (A) patient age and (B) body temperature at triage. Age was binned in 5 years intervals, and temperature in 1.5 °C degrees intervals.*

**Figure S4: Case fatality rate as a function of cycle threshold**

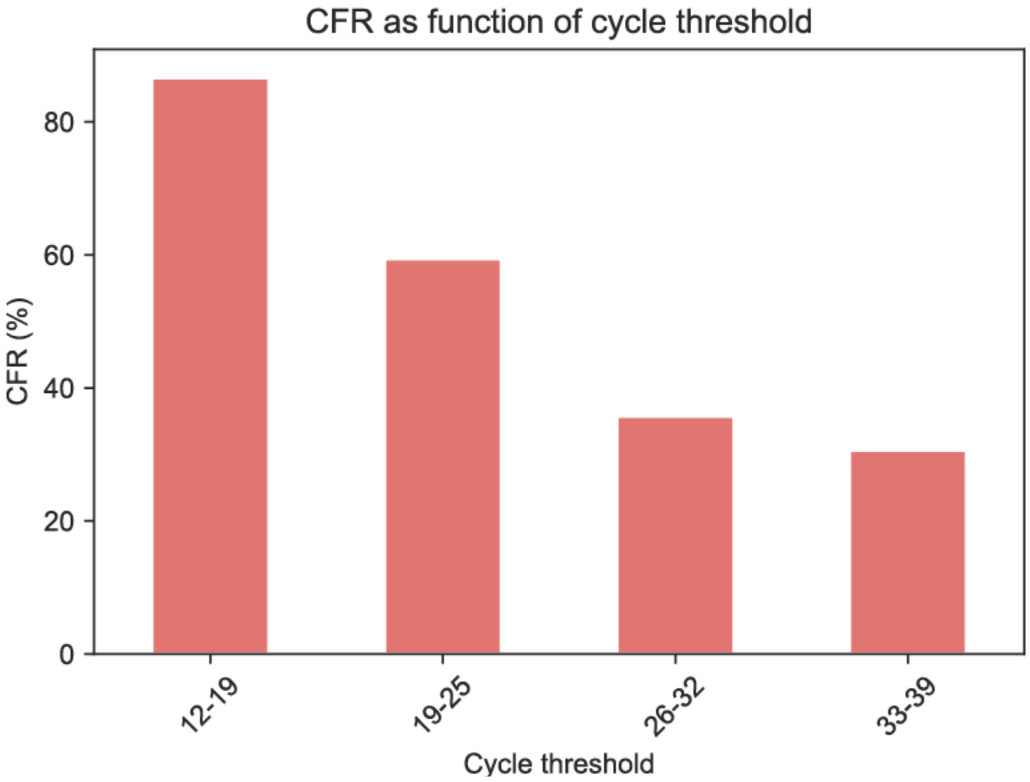


*Case Fatality Rate as a function of the measured RT-PCR cycle threshold values measured from the first or second blood draw.*

**Table S1: Model specifications**

**S1A: Parsimonious model**

| Term | Coefficient | P-value | Odds ratios | Risk ratios | Type | Knots |
|---|---|---|---|---|---|---|
| *Intercept* | 8.14656178 | 0.5795 | | | constant | |
| *Patient Age* | -0.0588981 | 0.0004 | | | RCS0 | |
| *Patient Age'* | 0.10325771 | <0.0001 | | | RCS1 | 5 30 58 |
| *Cycle Threshold* | -0.6919651 | 0.0013 | 0.42 | | linear | |
| *Body Temperature* | -0.2056551 | 0.6050 | | | RCS0 | |
| *Body Temperature'* | 0.75788984 | 0.2541 | | | RCS1 | 36.3 37.2 39.31 |
| *Jaundice* | 1.33911639 | 0.0360 | 3.82 | 1.04 | linear | |
| *Bleeding* | 1.02004343 | 0.0511 | 2.77 | 1.04 | linear | |
| *Dyspnea* | 0.17193284 | 0.5388 | 1.19 | 1.03 | linear | |
| *Dysphagia* | 0.14673348 | 0.5992 | 1.16 | 1.03 | linear | |
| *Referral Time* | 0.00119668 | 0.9759 | 1.01 | | linear | |
| *Cycle Threshold x Referral Time* | -0.0420558 | 0.2926 | | | product | |

**Formula:** 8.14656 - 0.0589 age + $4\times10^{-5}$ max(age - 5.0, 0)$^3$ – $7\times10^{-5}$ max(age - 30.0, 0)$^3$ + $3\times10^{-5}$ max(age - 58.0, 0)$^3$ - 0.69197 CT - 0.20566 temp + 0.08365 max(temp - 36.3, 0)$^3$ - 0.11933 max(temp - 37.2, 0)$^3$ + 0.03568 max(temp - 39.31, 0)$^3$ + 1.33912 jaundice + 1.02004 bleeding + 0.17193 dyspnea + 0.14673 dysphagia + 0.0012 RT - 0.04206 CT x RT

**S1B: Parsimonious model without body temperature**

| Term | Coefficient | P-value | Odds ratios | Risk ratios | Type | Knots |
|---|---|---|---|---|---|---|
| *Intercept* | 0.79342334 | 0.0120 | | | constant | |
| *Patient Age* | -0.0596321 | 0.0003 | | | RCS0 | |
| *Patient Age'* | 0.10436838 | <0.0001 | | | RCS1 | 5 30 58 |
| *Cycle Threshold* | -0.6944901 | 0.0011 | 0.42 | | linear | |
| *Jaundice* | 1.31695855 | 0.0373 | 3.72 | 1.04 | linear | |
| *Bleeding* | 0.92970184 | 0.0725 | 2.53 | 1.03 | linear | |
| *Dyspnea* | 0.16771205 | 0.5458 | 1.18 | 1.04 | linear | |
| *Dysphagia* | 0.08827503 | 0.7442 | 1.09 | 1.02 | linear | |
| *Referral Time* | -0.0023547 | 0.9515 | 1.00 | | linear | |
| *Cycle Threshold x Referral Time* | -0.04104 | 0.3011 | | | product | |

**Formula:** 0.79342 - 0.05963 age + $4x10^{-5}$ max(age - 5.0, $0)^3$ – $7x10^{-5}$ max(age - 30.0, $0)^3$ + 3e-05 max(age - 58.0, $0)^3$ - 0.69449 CT + 1.31696 jaundice + 0.9297 bleeding + 0.16771 dyspnea + 0.08828 dysphagia - 0.00235 RT - 0.04104 CR x RT

#### S1C: Clinical-only model

| Term | Coefficient | P-value | Odds ratios | Risk ratios | Type | Knots |
|---|---|---|---|---|---|---|
| Intercept | 3.02919786 | 0.8213 | | | constant | |
| Patient Age | -0.0527675 | 0.0004 | | | RCS0 | |
| Patient Age' | 0.08708779 | <0.0001 | | | RCS1 | 5 30 58 |
| Body Temperature | -0.0752819 | 0.8355 | | | RCS0 | |
| Body Temperature' | 0.57465845 | 0.3436 | | | RCS1 | 36.3 37.2 39.31 |
| Jaundice | 1.30197731 | 0.0241 | 3.68 | 1.03 | linear | |
| Bleeding | 1.16552489 | 0.0149 | 3.21 | 1.03 | linear | |
| Dyspnea | 0.16085186 | 0.5328 | 1.17 | 1.04 | linear | |
| Dysphagia | 0.27387136 | 0.2891 | 1.31 | 1.05 | linear | |
| Asthenia/weakness | 0.23194879 | 0.3249 | 1.26 | 1.16 | linear | |
| Diarrhea | 0.14197065 | 0.5163 | 1.15 | 1.07 | linear | |

**Formula:** 3.0292 - 0.05277 age + $3x10^{-5}$ max(age - 5.0, $0)^3$ - 6e-05 max(age - 30.0, $0)^3$ + $3x10^{-5}$ max(age - 58.0, $0)^3$ - 0.07528 temp + 0.06343 max(temp - 36.3, $0)^3$ - 0.09048 max(temp - 37.2, $0)^3$ + 0.02705 max(temp - 39.31, $0)^3$ + 1.30198 jaundice + 1.16552 bleeding + 0.16085 dyspnea + 0.27387 dysphagia + 0.23195 asthenia + 0.14197 diarrhea

#### S1D: Minimal model

| Term | Coefficient | P-value | Odds ratios | Risk ratios | Type | Knots |
|---|---|---|---|---|---|---|
| Intercept | 0.86215216 | 0.0042 | | | constant | |
| Patient Age | -0.0534333 | 0.0007 | | | RCS0 | |
| Patient Age' | 0.09478786 | <0.0001 | | | RCS1 | 5 30 58 |
| Cycle Threshold | -0.8694039 | <0.0001 | 0.34 | | linear | |

**Formula:** 0.86215 - 0.05343 age + $3x10^{-5}$ max(age - 5.0, $0)^3$ – $6x10^{-5}$ max(age - 30.0, $0)^3$ + $3x10^{-5}$ max(age - 58.0, $0)^3$ - 0.8694 CT

**S1E: Wellness scale parsimonious model**

| Term | Coefficient | P-value | Odd ratios | Risk ratios | Type | Knots |
|---|---|---|---|---|---|---|
| Intercept | 21.379251 | 0.2201 | | | constant | |
| Patient Age | -0.0754931 | 0.0014 | | | RCS0 | |
| Patient Age' | 0.13750982 | <0.0001 | | | RCS1 | 4 28 60 |
| Cycle Threshold | -0.5247369 | 0.0862 | 0.51 | | linear | |
| Body Temperature | -0.5798065 | 0.2196 | | | RCS0 | |
| Body Temperature' | 1.25110525 | 0.1077 | | | RCS1 | 36.2 37.2 39.29 |
| Wellness Scale | 0.49509775 | 0.0142 | 2.69 | | linear | |
| Referral Time | -0.0930255 | 0.1502 | 0.39 | | linear | |
| Cycle Threshold x Referral Time | -0.106844 | 0.2122 | | | product | |

**Formula:** $21.37925 - 0.07549\ \text{age} + 4\text{x}10^{-5} \max(\text{age} - 4.0, 0)^3 - 8\text{x}10^{-5} \max(\text{age} - 28.0, 0)^3 + 3\text{x}10^{-5} \max(\text{age} - 60.0, 0)^3 - 0.52474\ \text{CT} - 0.57981\ \text{temp} + 0.13103 \max(\text{temp} - 36.2, 0)^3 - 0.19373 \max(\text{temp} - 37.2, 0)^3 + 0.06269 \max(\text{temp} - 39.29, 0)^3 + 0.4951\ \text{WS} - 0.09303\ \text{RT} - 0.10684\ \text{CT x RT}$

**S1F: Wellness scale parsimonious without body temperature model**

| Term | Coefficient | P-value | Odds ratios | Risk ratios | Type | Knots |
|---|---|---|---|---|---|---|
| Intercept | 0.17095348 | 0.8039 | | | constant | |
| Patient Age | -0.0749107 | 0.0012 | | | RCS0 | |
| Patient Age' | 0.1384085 | <0.0001 | | | RCS1 | 4 28 60 |
| Cycle Threshold | -0.5033504 | 0.0930 | 0.53 | | linear | |
| Wellness Scale | 0.50788889 | 0.0108 | 2.76 | | linear | |
| Referral Time | -0.0968393 | 0.1256 | 0.38 | | linear | |
| Cycle Threshold x Referral Time | -0.1070374 | 0.2070 | | | product | |

**Formula:** $0.17095 - 0.07491\ \text{age} + 4\text{x}10^{-5} \max(\text{age} - 4.0, 0)^3 - 8\text{x}10^{-5} \max(\text{age} - 28.0, 0)^3 + 3\text{x}10^{-5} \max(\text{age} - 60.0, 0)^3 - 0.50335\ \text{CT} + 0.50789\ \text{WS} - 0.09684\ \text{RT} - 0.10704\ \text{CT x RT}$

**S1G: Wellness scale clinical-only model**

| Term | Coefficient | P-value | Odds ratios | Risk ratios | Type | Knots |
|---|---|---|---|---|---|---|
| Intercept | 13.0101569 | 0.3890 | | | constant | |
| Patient Age | -0.0691352 | 0.0006 | | | RCS0 | |
| Patient Age' | 0.123125 | <0.0001 | | | RCS1 | 4 28 60 |
| Body Temperature | -0.3840269 | 0.3494 | | | RCS0 | |
| Body Temperature' | 0.99239011 | 0.1543 | | | RCS1 | 36.2 37.2 39.29 |
| Wellness Scale | 0.74786034 | <0.0001 | 2.11 | | linear | |

**Formula:** 13.01016 - 0.06914 age + $4x10^{-5}$ max(age - 4.0, 0)$^3$ - $7x10^{-5}$ max(age - 28.0, 0)$^3$ + $3x10^{-5}$ max(age - 60.0, 0)$^3$ - 0.38403 temp + 0.10394 max(temp - 36.2, 0)$^3$ - 0.15367 max(temp - 37.2, 0)$^3$ + 0.04973 max(temp - 39.29, 0)$^3$ + 0.74786 WS

**S1H: Wellness scale minimal model**

| Term | Coefficient | P-value | | Type | Knots |
|---|---|---|---|---|---|
| Intercept | 0.17627324 | 0.7957 | | constant | |
| Patient Age | -0.0826796 | 0.0002 | | RCS0 | |
| Patient Age' | 0.14654649 | <0.0001 | | RCS1 | 4 28 60 |
| Cycle Threshold | -0.7917185 | 0.0002 | 0.38 | linear | |
| Wellness Scale | 0.43591077 | 0.0198 | 2.39 | linear | |

**Formula:** 0.17627 - 0.08268 age + $5x10^{-5}$ max(age - 4.0, 0)$^3$ - $8x10^{-5}$ max(age - 28.0, 0)$^3$ + $4x10^{-5}$ max(age - 60.0, 0)$^3$ - 0.79172 CT + 0.43591 WS

*Tables describing all the models constructed in this paper: (A) parsimonious, (B) parsimonious without body temperature, (C) clinical-only, (D) minimal, (E) wellness scale parsimonious, (F) wellness scale parsimonious without body temperature, (G) wellness scale clinical-only, and (H) wellness scale minimal. The tables provide the model coefficients for each term, their P-values, odds ratios for the linear terms (caused by the presence of a sign/symptom or an IQR change in continuous variables), risk ratios (estimated from the odds ratios only for categorical variables), type of term (constant, linear, restricted cubic spline or RCS, and product), and the knots of the non-linear RCS term (RCS1). The formula of the logit(p) function for each model is provided as well.*

**Table S2: Performance comparison between CT day 1 and 2**

| *Disposition* | *Age* | *CT1* | *CT2* | *Temperature* | *Jaundice* | *Bleeding* | *Dyspnea* | *Dysphagia* | *Referral Time* | *Score1* | *Score1* |
|---|---|---|---|---|---|---|---|---|---|---|---|
| *Survived* | 4 | 0.885 | 0.885 | 36.7 | No | No | No | No | 1 | 0.431 | 0.431 |
| *Survived* | 8 | -0.202 | 0.303 | 39.3 | No | No | No | No | 2 | 0.712 | 0.626 |
| *Died* | 50 | -0.163 | -0.027 | 38.6 | No | No | No | Yes | 1 | 0.734 | 0.714 |
| *Died* | 40 | 0.594 | 0.652 | 40 | No | No | No | No | 1 | 0.566 | 0.555 |
| *Died* | 28 | -0.454 | -0.512 | 38.9 | No | No | Yes | No | 3 | 0.593 | 0.604 |
| *Died* | <1 | 0.691 | 0.691 | 38.2 | No | No | No | No | 1 | 0.553 | 0.553 |
| *Survived* | 38 | 0.400 | 2.282 | 36.6 | No | No | No | No | 3 | 0.342 | 0.100 |
| *Survived* | 27 | -0.493 | -0.434 | 36.7 | No | No | No | No | 2 | 0.448 | 0.437 |
| *Died* | 16 | -0.919 | 2.204 | 36.3 | No | Yes | Yes | No | 7 | 0.869 | 0.234 |

*The table shows the information of the patients with CT data for both days 1 and 2, as required to evaluate the parsimonious model (age, CT, body temperature, jaundice, bleeding, dyspnea, dysphagia, and referral time). The CT values are site-normalized (by subtracting the mean and dividing by the standard deviation of the CT values either from the Sierra Leonean or Liberian ETUs, depending on where the patient was treated). The scores 1 and 2 are the predictions from the parsimonious when using CT1 or CT2 as the input for CT. The classification threshold is 0.5, so the prediction of the model is death if the score is greater than 0.5.*

**Table S3: Performance indices of wellness models without imputation**

| | *WS parsimonious (95% CI)* | *WS parsimonious w/out temp. (95% CI)* | *WS clinical-only (95% CI)* | *WS minimal (95% CI)* |
|---|---|---|---|---|
| *AUC* | 0.80 (0.74, 0.84) | 0.80 (0.75, 0.85) | 0.74 (0.67, 0.79) | 0.80 (0.74, 0.85) |
| $R^2$ | 0.31 (0.24, 0.38) | 0.30 (0.23, 0.37) | 0.20 (0.15, 0.26) | 0.25 (0.19, 0.32) |
| *Brier* | 0.19 (0.13, 0.24) | 0.18 (0.13, 0.24) | 0.21 (0.15, 0.27) | 0.18 (0.13, 0.24) |
| *Accuracy* | 0.73 (0.66, 0.79) | 0.72 (0.66, 0.78) | 0.68 (0.60, 0.74) | 0.72 (0.65, 0.77) |
| *Sensitivity* | 0.81 (0.75, 0.85) | 0.81 (0.76, 0.86) | 0.78 (0.72, 0.84) | 0.81 (0.75, 0.85) |
| *Specificity* | 0.64 (0.56, 0.71) | 0.61 (0.53, 0.68) | 0.55 (0.47, 0.63) | 0.60 (0.52, 0.67) |

*Performance indices of the wellness scale models obtained without applying multiple imputation on the wellness scale.*

**PCR Lab Notes**

Liberia

For both the Bong and Margibi ETUs, laboratory diagnosis of Ebola virus disease was performed at the United States Naval Medical Research Center (NMRC) Mobile Laboratory in Bong County, Liberia, with the 1-step quantitative Ebola Zaire real-time reverse transcriptase–polymerase chain reaction (TaqMan) assay (Naval Medical Research Center, Frederick, MD). Briefly, Qiagen buffer AVL/ethanol-inactivated blood samples were extracted with QIAamp Viral RNA Mini Kit. Extracted ribonucleic acid was tested for 2 Ebola virus disease gene targets (EBOV Zaire locus and minor groove binding locus), using the Applied Biosystems StepOnePlus instrument. A sample was confirmed to be positive for Ebola virus disease if both targets were detected.

Sierra Leone

The Public Health England (PHE) Port Loko lab (serving Lunsar ETU) switched Ebola real-time PCR assay from commercial Altona –widely used by other "mobile" labs in West Africa – to the more sensitive and robust in-house "Trombley" on 2/3/15. The Public Health England Makeni lab (serving Makeni ETU) had made the switch on 2/4/15. Both assays tested a single EVD gene target. The Nigerian Lab (serving Kambia ETU) used the Altona assay the entire time.

Briefly, the process used was that the plasma sample (80ul) was inactivated in Qiagen AVL (320 µl) /10mins, then heat-treated for 15 min at 60C, prior to extraction using a Qiagen EZ1 virus kit V2.0. The eluted nucleic acid was analysed using a one-step real time PCR directed against the EBOV Zaire nucleoprotein region (based on the PCR by Trombley et al [3], with some modifications including multiplexing with an Internal control [1, 2]), and performed on a Smartcycler PCR machine.

The NRMC lab assays and processing methodologies are fairly similar to the PHE labs. The NRMC use two targets, including the EBOV Zaire target and the minor groove binding target, whereas the PHE lab used only the EBOV Zaire target. The other main difference was the PHE extraction machine was automated whereas the NRMC was a manual methodology, but both would be expected to generate similar results.

More specific details are in the papers below.

1. **Weller SA, Bailey D, Matthews S, Lumley S, Sweed A, Ready D, Eltringham G, Richards J, Vipond R, Lukaszewski R, Payne PM, Aarons E, Simpson AJ, Hutley EJ, Brooks T.** 2016. Evaluation of the Biofire FilmArray BioThreat-E Test (v2.5) for Rapid Identification of Ebola Virus Disease in Heat-Treated Blood Samples Obtained in Sierra Leone and the United Kingdom. J Clin Microbiol **54:**114-119.
2. **Broadhurst MJ, Kelly JD, Miller A, Semper A, Bailey D, Groppelli E, Simpson A, Brooks T, Hula S, Nyoni W, Sankoh AB, Kanu S, Jalloh A, Ton Q, Sarchet N, George P, Perkins MD, Wonderly B, Murray M, Pollock NR.** 2015.

ReEBOV Antigen Rapid Test kit for point-of-care and laboratory-based testing for Ebola virus disease: a field validation study. Lancet **386:**867-874.
3. **Trombley AR, Wachter L, Garrison J, Buckley-Beason VA, Jahrling J, Hensley LE, Schoepp RJ, Norwood DA, Goba A, Fair JN, Kulesh DA.** 2010. Comprehensive panel of real-time TaqMan polymerase chain reaction assays for detection and absolute quantification of filoviruses, arenaviruses, and New World hantaviruses. Am J Trop Med Hyg **82:**954-960.
